# Resource limitation reveals a twofold benefit of eusociality

**DOI:** 10.1101/053108

**Authors:** Emanuel A. Fronhofer, Jürgen Liebig, Oliver Mitesser, Hans Joachim Poethke

## Abstract

Explaining the evolution and maintenance of cooperative breeding or eusociality remains a challenge. Surprisingly, fundamental ecological factors, specifically competition for limited resources and resource variance, are frequently ignored in models of animal sociality. We here develop a mathematical model that includes density-dependent population growth and quantify the influence of cooperative foraging on resource use efficiency. We derive optimal resource sharing strategies, ranging from egalitarian to cooperatively breeding and eusocial groups. We find that, while egalitarian resource sharing is a risk-reducing foraging strategy, eusociality yields additional benefits: like egalitarian strategies, eusocial groups can reduce their members’ starvation risk by reducing resource variance. Additionally, eusocial groups increase their reproductive output by increasing intra-group variance in resources allocated to reproduction. This allows reproduction even when resources are so scarce that solitary animals would not be able to reproduce. In a majority of environmental situations and life-histories, this twofold benefit of eusociality increased resource use efficiency and led to supersaturation, that is, to a strong increase in carrying capacity. Supersaturation provides indirect benefits to group members even for low intra-group relatedness and represents one potential explanation for the evolution and maintenance of eusociality and cooperative breeding.

## Introduction

The evolution and maintenance of cooperative behavior in animals has been a topic of ongoing interest since the days of Darwin. A number of possible factors favoring cooperation have been proposed (reviewed in Krause and Ruxton, 2002; Nowak, 2006; Lehmann and Keller, 2006). Generally, direct benefits of cooperation lead to increased own reproduction, while indirect benefits are received through increased reproduction of relatives (Hamilton, 1964a,b). Thus, relatedness is a major factor for the evolution of cooperation with indirect benefits, such as cooperative breeding or eusociality. However, the origins of these social systems have been associated with biparental families (Hughes et al., 2008; Cornwallis et al., 2010; Boomsma, 2013) which results in offspring being equally related to own offspring and to brothers or sisters (e.g., Bourke and Franks, 1995). In this context, whether individuals should favor own offspring production or the raising of siblings depends less on relatedness, which is symmetrical, but more on ecological factors and constraints (see e.g, Avila and Fromhage, 2015), such as food availability, that make either helping or solitary breeding more successful (West et al., 2007).

Although food is a major determinant of an individual’s survival and reproduction, the role of resource availability and its variability, which are important ecological factors (Hatchwell and Komdeur, 2000), is often underestimated. Furthermore, the fact that resource availability is not a fixed environmental factor, but an emergent quantity which depends not only on the environment but also on population size, is only rarely acknowledged (but see Pen and Weissing, 2000; López-Sepulcre and Kokko, 2005). In the following model, we therefore focus on the interaction between resource acquisition and allocation on the one hand and resource availability on the other and demonstrate the impact of these ecological factors on group formation.

Foraging decisions are affected by the mean amount of resources as well as by their variance, as suggested by risk-sensitive foraging theory (reviewed in Bateson, 2002; Bednekoff, 1996; Kacelnik and Bateson, 1996; Smallwood, 1996). In this context, group formation has traditionally been seen as a risk-averse, i.e. variance reducing, mechanism (Caraco, 1981; Clark and Mangel, 1986; Wenzel and Pickering, 1991; Caraco et al., 1995; Uetz, 1996; Uetz and Hieber, 1997). The simple idea behind these models is that foraging success may vary: individuals may find resources that are too large to fully utilize, or alternatively, they may not find any resources, which leads to certain death (Fig. 1 B). Foraging with subsequent egalitarian resource sharing allows animals to dampen such environmental variation (Fig. 1 A), as all group members will receive an intermediate amount of resources which guarantees positive long-term fitness (given a sufficient level of resource variance; see also Fronhofer et al., 2011a).

**Figure 1:**
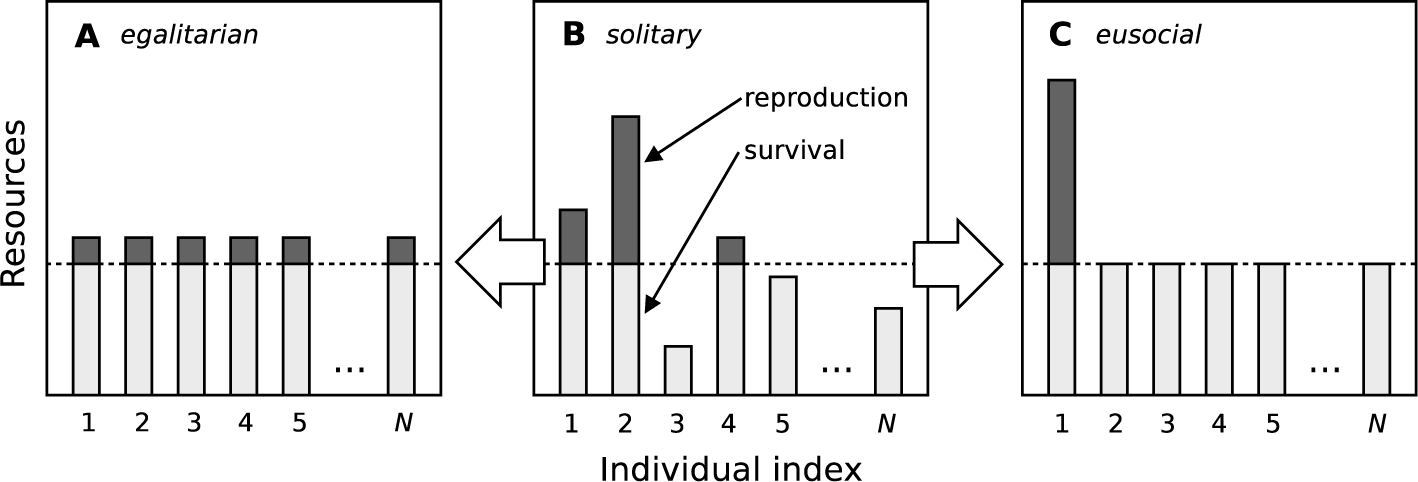
Schematic comparison of different modes of resource distribution in groups of animals: Individuals collect resources individually and vary in their success. Solitary individuals (B) thus differ in the amount of resources individuals may use for survival (light gray) and reproduction (dark gray). Some individuals (3, 5, and *N*) will die of starvation. In egalitarian groups (A) resources are evenly shared after solitary foraging (B). All individuals can survive and receive a small amount for reproduction. In eusocial groups (C) individuals receive sufficient resources for survival and channel all remaining resources to the reproductive dominant individual (here individual 1).

Yet, as Poethke and Liebig (2008) point out, group formation is not necessarily a variance-reducing mechanism. It may be seen as an important means of variance manipulation in general: whether variance in resource availability is reduced or increased depends on the degree of reproductive division of labor. While egalitarian resource allocation decreases intra-group variance as explained above (Fig. 1 A), skewed resource allocation, by contrast, increases variance (Fig. 1 C). If resource availability and variance are low solitary foragers may collect more food than needed for survival, but not enough to reproduce. If individuals forage cooperatively, subsequently pool the surplus of resources not needed for survival and then redirect this surplus towards one (or a few) individual(s), individuals in groups will survive and specific group members have a chance to reproduce. This clearly increases fitness, either through direct fitness benefits (for the reproductive individual) or indirect fitness benefits (due to intra-group relatedness).

Thus, in principle, two fundamentally different types of cooperative animal groups exist (see Tab. 1). On the one hand, individuals forming egalitarian groups forage and subsequently share the pooled resources so that every group member receives roughly the same amount of food (examples include lions and social spiders: Packer et al., 2001; Whitehouse and Lubin, 2005). Of course, this is a rare situation at one end of a continuum of different degrees of reproductive division of labor (i.e. skew, for a review see Reeve and Keller, 2001). On the other hand, one can find animal societies in which just one individual receives all the resources for reproduction while the other members of the group only obtain a share necessary for their survival (e.g., eusocial insects or mole-rats; Wilson, 1971; Clutton-Brock et al., 2009, such groups are henceforth termed “eusocial”; in the literature one may also find the term “despotic” which would be equivalent here). Evidently, as Sherman et al. (1995) point out, other degrees of reproductive division of labor in between these two extremes are possible and often encountered (for numerous examples from insect societies alone see Wilson, 1971; Hölldobler and Wilson, 1990, 2009; Costa, 2006).

**Table 1:** Use and intra-group distribution of resources in groups of different social organization. While egalitarian societies pool the (individually collected) resources and equally share them between the members of the group, eusocial societies channel all resources not needed for survival to one (or a few) reproductive individuals. Subordinates will usually keep (or get from the pool) sufficient resources for survival and only deliver the remaining amount to the reproductive individual. The reproductive individual may—in a strictly despotic society—even decide whether individuals are fed or the resources are completely invested into reproduction. We do not explicitly consider the latter case here, as further analyses suggest that this type of group is even more efficient. Our results are therefore conservative.

| **type of group** | **use of resources** | **survival** | **reproduction** |
| --- | --- | --- | --- |
| solitary | individually | — | — |
| egalitarian | pooled | egalitarian | egalitarian |
| eusocial | pooled | egalitarian | despotic |
| strictly despotic | pooled | despotic | despotic |

As a consequence of these considerations, previous work by Poethke and Liebig (2008) suggests that egalitarian societies, as a risk-reducing foraging strategy, should be favored in environments with high resource variance and eusocial animal societies in habitats with low resource variance, since this group structure increases inter-individual variance. Yet, in nature, egalitarian animal societies are only rarely found (Packer et al., 2001). We assume that this discrepancy between model predictions and empirical observations stems from the fact that previous theoretical work does not take into account density-dependent population growth, i.e. the interaction between population size and food availability. However, density-dependence has been shown to be of high relevance in the context of risk-sensitive foraging in general (Fronhofer et al., 2011b). Therefore, we here develop a mathematical model that accounts for the influence of foraging and resource allocation on population size and resource availability, that is, density-dependence. We compare cooperative breeders or eusocial groups, i.e. groups in which only one animal is allowed to reproduce in our model, with egalitarian societies and solitary strategies (i.e. groups of size one), and present one possible explanation for the dominance of eusocial groups in nature.

## Model description and numerical results

### Resource availability

We assume stochastic foraging, that is, individual foraging success follows a random distribution and the per capita probability of collecting an amount x of resources during one time step is given by a probability density function 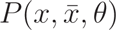. We assume that individuals collect resource items of limited size. Thus, variance in foraging success is determined by the mean resource item size *θ*. As the amount of resources collected should be non-negative and continuous, the distribution of resources is easily described by a Gamma distribution:

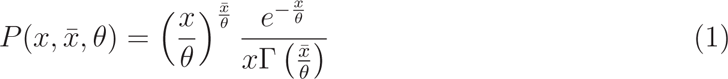

with mean 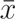, scale parameter *θ*, and the gamma function Г (Andrews et al., 2001). The variance of acquired resources is 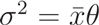 and the coefficient of variation 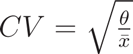. For a constant mean amount of food 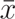 collected by an individual an increase in item size will necessarily be accompanied by an increase in the variance of the amount of resources collected (Fig. 2 A). In the following, we will therefore use mean resource item size *θ* as a proxy for environmental variance.

**Figure 2:**
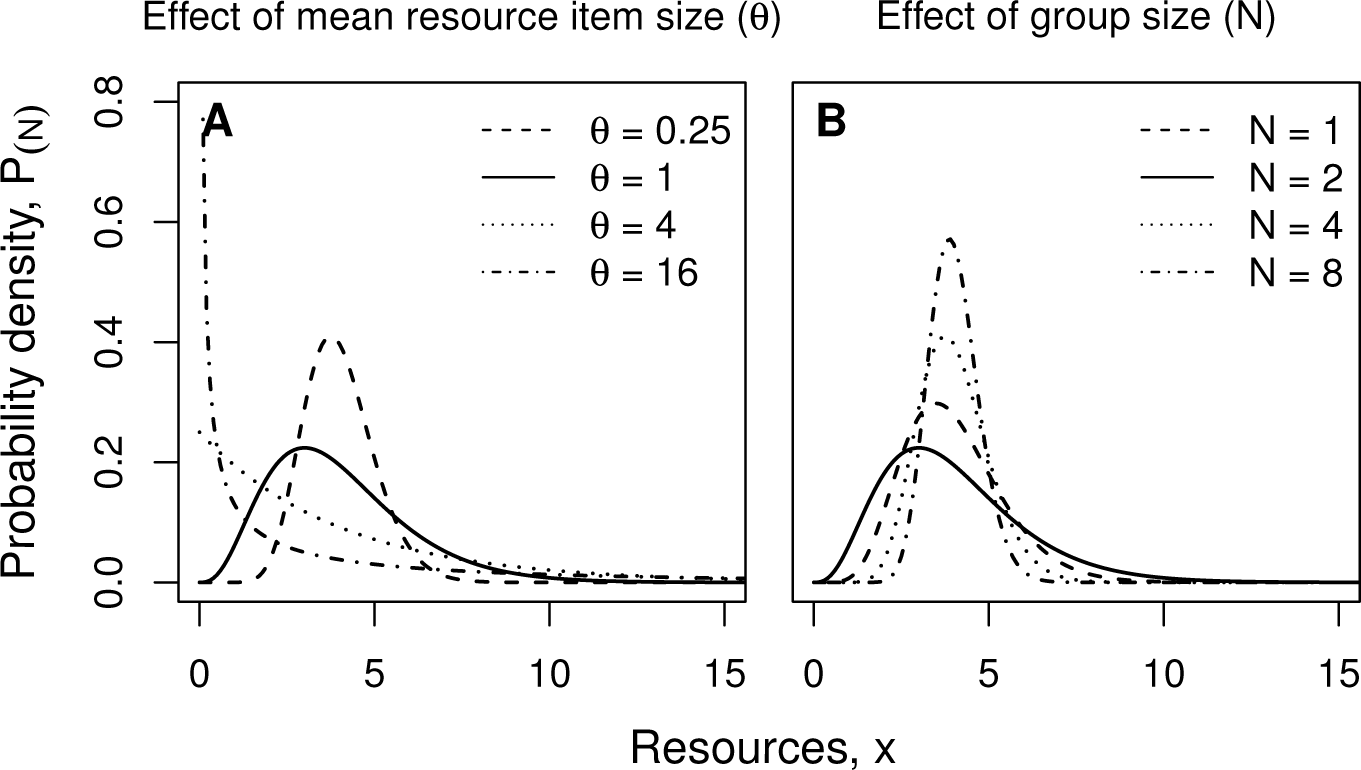
Distribution of resources (*x*) available to an individual. Influence of mean resource item size *θ* (A) and group size *N* (B) on the distribution of resources x available to an individual as a solitary 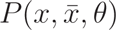 (Eqn. 1) respectively 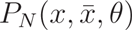 in a group of size *N* (Eqn. 2). (A): 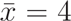; *N* =1; *θ* = 0.25 (dashed line), *θ* = 1 (solid line), *θ* = 4 (dotted line), *θ* = 16 (dotdashed line). (B): 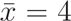; *θ* =1; *N* =1 (solid line), *N* = 2 (dashed line), *N* = 4 (dotted line), *N* = 8 (dotdashed line).

For both, individuals in egalitarian as well as individuals in eusocial groups, we assume that foraged resources are pooled and subsequently allocated to survival and reproduction (for an overview see Tab. 1). This process of resource pooling modifies the resource distribution (Fig. 2 B) as detailed in Poethke and Liebig (2008) and Fronhofer et al. (2011a). Basically, variance in resources decreases with 1/*N*. Thus, the amount of resources available per individual in a group of size *N* follows a modified Gamma distribution with reduced variance:

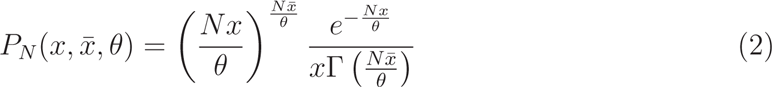

### Fertility and mortality

We assume that individual mortality *M* is a function of the amount of resources *x_s_* allocated to survival and, as a simplification, we employ a step function. That is, we assume that an animal dies if it receives less resources than a certain threshold resource value (*o_M_*) and survives with a certain probability if it receives more (Fig. 3):

**Figure 3:**
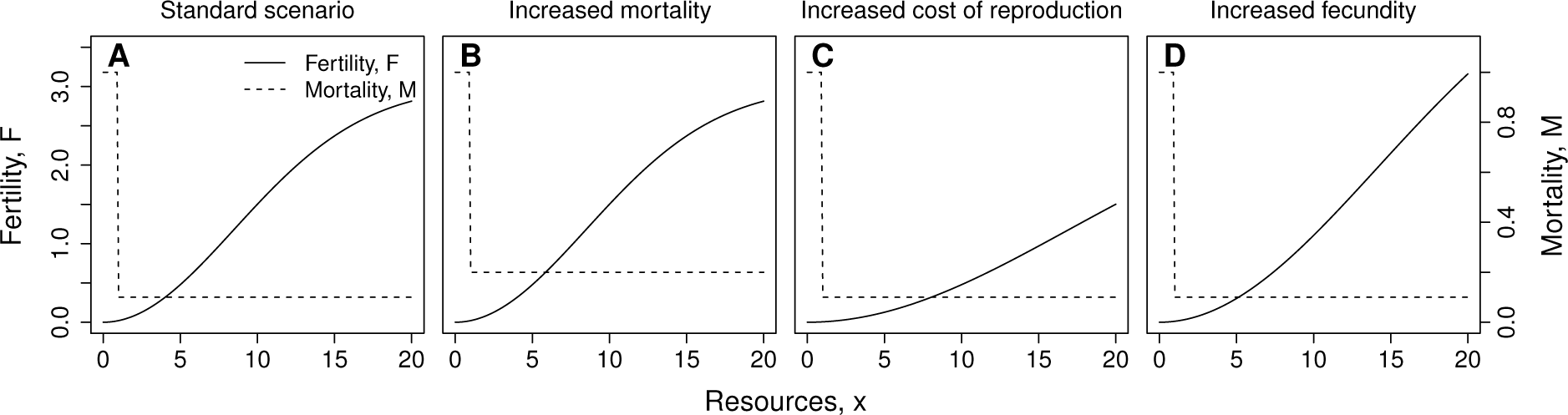
Influence of model parameters on fertility function (*F*(*x* = *x*_*r*_); solid line) and mortality function (*M*(*x* = *x*_*s*_); dashed line) for four exemplary parameter combinations. (A): standard parameter set (*F*_*max*_ = 3; *c*_0_ = 4; *M*_*b*_ = 0.1; *o*_*M*_ = 1); (B): increased mortality (*F*_*max*_ = 3; *c*_0_ = 4; *M*_*b*_ = 0.2; *o*_*M*_ = 1); (C): increased cost of reproduction (*F*_*max*_ = 3; *c*_*0*_ = 8; *M*_*b*_ = 0.1; *o*_*M*_ = 1); (D): increased fecundity (*F*_*max*_ = 5; *c*_0_ = 4; *M*_*b*_ = 0.1; *o*_*M*_ = 1).

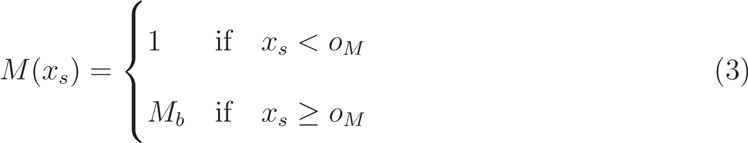

with the resource independent baseline mortality *M_b_,* resulting from predation or disease, for instance. We have analyzed the influence of including a sigmoid function for mortality and could show that this does not change our results qualitatively. More generally, the consequences of the form of the fitness function (mortality and fertility, see below) is discussed in Fronhofer et al. (2011b) in the context of risk-sensitive foraging.

Pooled resources are first allocated to survival of group members until all individuals have received the amount *x_s_* = *o_M_* preventing death from starvation. Individuals die if there are not sufficient resources available. We thus get for the per capita mortality in a group of size *N*:

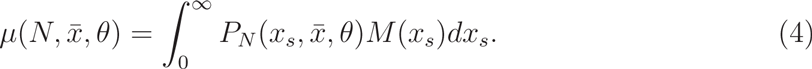

All remaining resources are allocated to reproduction, either by giving them to a single reproductively dominant individual (eusocial groups) or by equally sharing them between all members of the group (egalitarian groups; see Tab. 1). Given that the benefits of resource sharing will thus impact diferent group members diferently, we will investigate the relative success of individuals which do not stay in the group but attempt independent breeding below.

In general, reproduction *F* is a function of the resources available per capita *x*. As fertility is not unlimited, the functional relationship between fertility and the resources remaining after consumption for survival *x_r_* = max(0, *x* – *o_M_*) allocated to reproduction can be assumed to follow a sigmoid shape (see Fronhofer et al., 2011a,b, and Fig. 3):

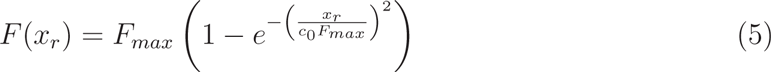

where *F_max_* determines fecundity, that is, the maximal value the reproduction function can take. For low values of *x_r_* the steepness of the fertility function is determined by 1 /*c*_0_. Therefore, *c*_0_ can be interpreted as the cost of reproduction. For an overview of parameter combinations under consideration see Tab. 2.

**Table 2:** Model parameters, meaning and tested values. Note that fecundity (*F_ma_*_*x*_) is a net rate, i.e. for solitaries *F_max_* = 5 leads to a quintupling of population size.

| parameter | values | meaning |
| --- | --- | --- |
| $F_{max}$ | [3, 5] | fecundity, i.e. maximal number of offspring |
| $c_0$ | [4, 8] | costs of offspring production |
| $\theta$ | [0, 32] | environmental variance (mean item size) |
| $M_b$ | [0.1, 0.2] | baseline mortality (resource independent) |
| $o_M$ | 1.0 | minimum amount of resources needed for survival |

Using Eqn. 5 we may calculate the per capita natality for individuals in groups of size *N* as a function of group size. For individuals in egalitarian groups one obtains

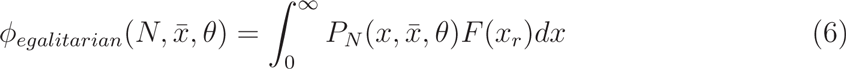

and for individuals in eusocial groups with only one reproductive individual

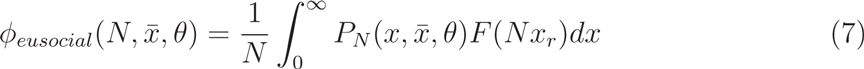

with *x_r_ =* max(0, *x – o_m_*).

Populations of individuals that compete for limited resources (density-dependent population growth) reach their stationary size (“carrying capacity”) when mortality balances natality We may thus formulate the equilibrium condition for populations consisting of individuals in groups of size *N* as

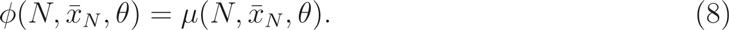

As both, per capita natality (Eqns. 6 and 7) and mortality (Eqn. 4), are functions of the distribution of resources acquired by individuals (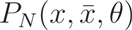, Eqn. 2), this yields an implicit relation that allows us to calculate – as a function of group size *N* and resource item size *θ* – the minimal mean amount of resources 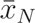 per individual needed to balance reproduction and mortality (Fig. 4).

**Figure 4:**
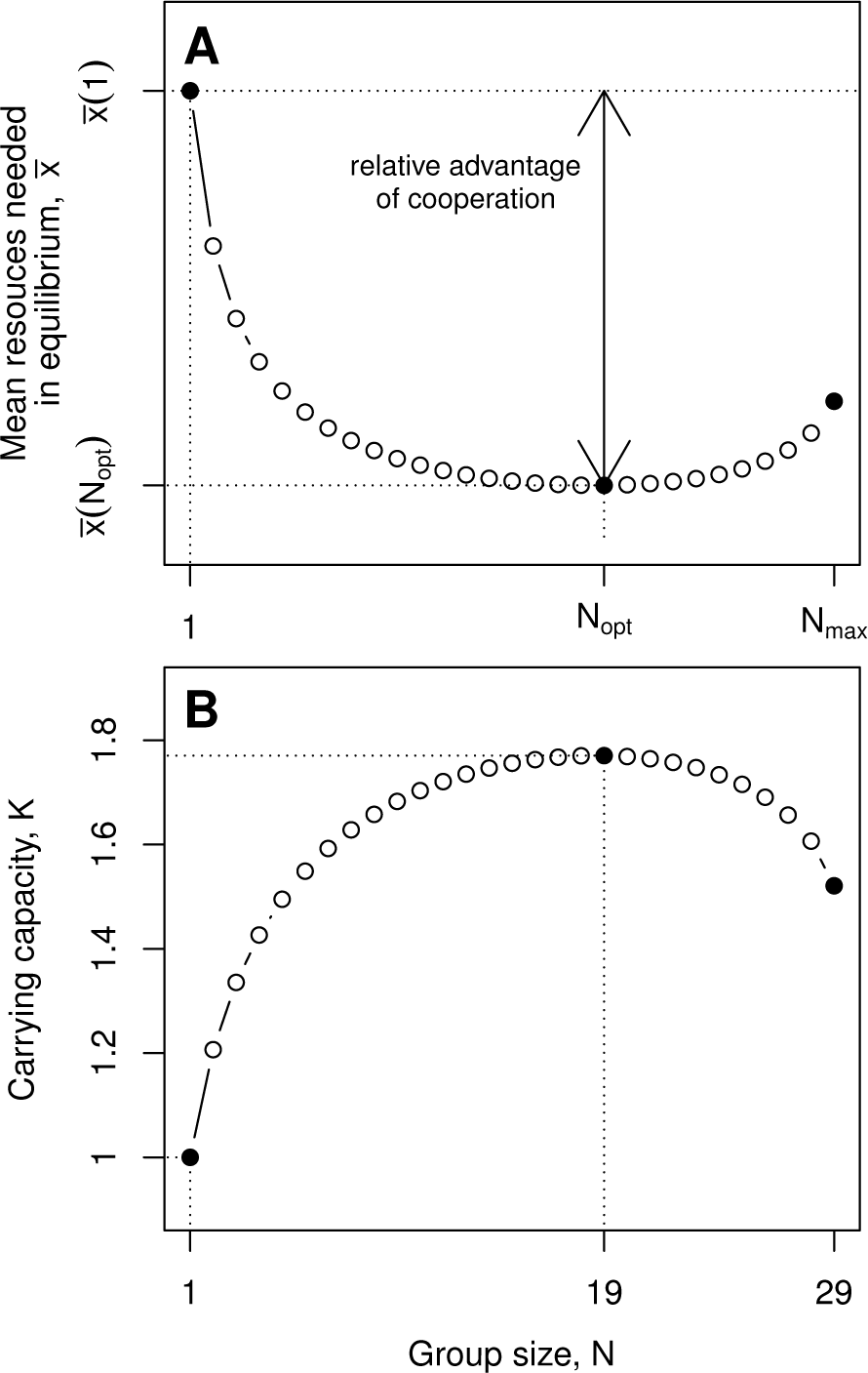
Influence of group size on (A) mean amount of resources needed to balance mortality and fertility in eusocial groups and (B) corresponding carrying capacity. Carrying capacity is shown relative to the carrying capacity of the solitary strategy. Numerical solution of Eqn. 8 for *F_max_* = 3, *c*_0_ = 4, *M_b_* = 0.1, *θ* = 5 and *o_M_* = 1.

In population equilibrium, evolution will minimize resource requirement (i.e., efficiency; respectively maximize carrying capacity; see among others MacArthur, 1962; MacArthur and Wilson, 1967; Boyce, 1984; Lande et al., 2009). A minimization of 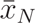 thus allows to determine the optimal group size *N_opt_* (Fronhofer et al., 2011a,b), i.e. the group size that maximizes carrying capacity (Fig. 4).

### Optimal group sizes and minimum resource requirements

As Eqn. 8 cannot be solved analytically, we approximated the results numerically. Fig. 5 gives the resulting mean amount of resources needed in population equilibrium and the resulting optimal group sizes for a broad range of environmental variance (0.1 ≤ *θ* ≤ 16). For a mean amount of resources collected per individual of approximately 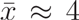, this corresponds to a coefficient of variation ranging from *CV* ≈ 0.16 to *CV* ≈ 2 (see Eqn. 1 and Fig. 2). The results (Fig. 5) clearly demonstrate the qualitative difference between egalitarian and eusocial group formation: While eusocial groups are advantageous at low resource variation, egalitarian groups become profitable when there is high variation in individual foraging success. This result confirms the general predictions of Poethke and Liebig (2008) who analyze the reproductive success of dyads and show the severe influence of variation in individual foraging success on the benefit of group formation. Poethke and Liebig (2008) state that “resource sharing in groups is a general mechanism of variance reduction while reproductive skew on the other hand allows increasing inter-individual variation in the amount of resources available for reproduction”.

**Figure 5:**
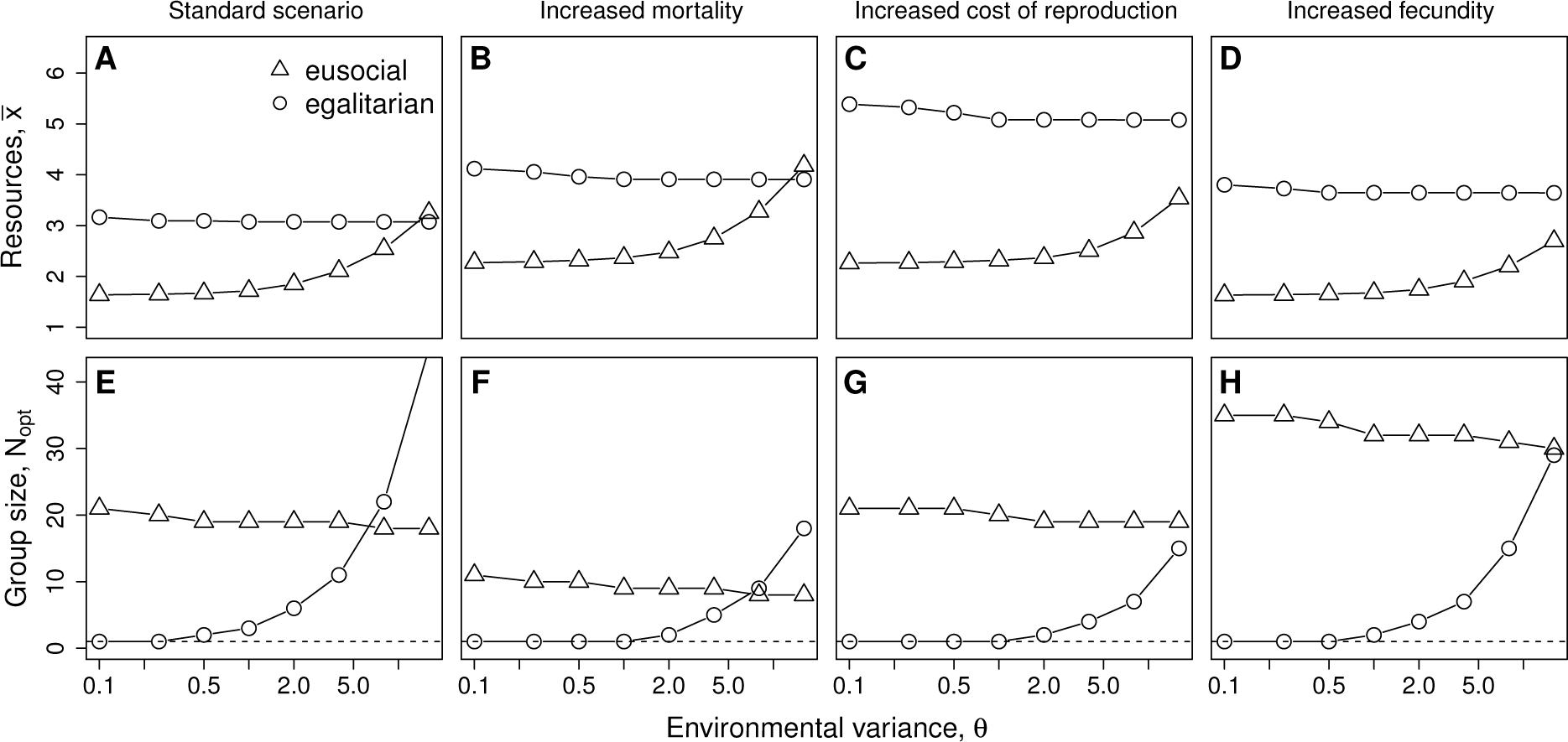
Influence of environmental variance (*θ*) on mean amount of resources needed (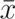, upper row) and optimal group size (*N_opt_*, lower row) for four exemplary parameter combinations (see Fig. 3). (A, E): standard parameter set (*F_max_* = 3; *c*_0_ = 4; *M_b_* = 0.1; *o_M_* = 1); (B, F): increased mortality (*F_max_* = 3; *c*_0_ = 4; *M_b_* = 0.2; *o_M_* = 1); (C, G): increased cost of reproduction (*F_max_* = 3; *c*_0_ = 8; *M_b_* = 0.1; *o_M_* = 1); (D, H): increased fecundity (*F_max_* = 5; *c*_0_ = 4; *M_b_* = 0.1; *o_M_* = 1). Circles give the results for egalitarian, triangles those for eusocial groups. Note the logarithmic x-axis.

#### Consequences of resource variance and the twofold benefit of eusociality

However, a closer look at our results reveals that reproductive skew in eusocial groups experiencing density-dependence does not simply increase inter-individual variance in the amount of resources available for reproduction. We may actually distinguish three fundamentally different situations: Firstly, for very low values of environmental variance (*θ* ≤ 1) a reduction of variance would actually decrease the expected fitness of individuals. This is due to the fact, that intra-specific competition necessarily results in low food availability and—as the fertility function is convex for low food availability (Fig. 3)—Jensen’s inequality (Ruel and Ayres, 1999) predicts a decrease in mean reproductive success for decreased variance in food availability. Thus, solitary strategies will outcompete egalitarian groups (i.e. optimal group sizes of egalitarian groups become *N_opt_* = 1; see Figs. 5 E, F, G, and H) in this area of parameter space. Eusocial groups, on the other hand, can increase inter-individual variance in food availability and will consequently out-compete solitary strategies (Figs. 5 A, B, C, and D).

Secondly, for intermediate values of environmental variation (1 < *θ* ≤ 10) variance reduction is obviously beneficial, which can be seen from the increase in optimal group sizes of egalitarian groups (Figs. 5 E, F, G, and H). Nevertheless, eusocial groups perform better than egalitarian groups in this range of environmental variances (*θ*). This is readily explained by the differential effect of variance in resource supply on fertility on the one hand and mortality on the other. While for this range of *θ* a reduction of variance still reduces mean reproductive success, it also reduces the mortality risk of individuals and, as long as the latter effect dominates, it pays off to reduce variance. However, eusocial groups may reduce variance in food supply for survival and, at the same time, increase variance in resources availability for reproduction. This twofold benefit of eusocial group formation explains the success of eusocial groups under a wide range of intermediate variance in foraging success.

Thirdly, if individual variance in foraging success is increased even further (*θ* ≥ 16), an increase of inter-individual variance in the amount of resources available for reproduction by channeling all resources to a reproductive dominant individual is no longer necessary and variance reduction by forming egalitarian groups may become the superior strategy depending on the other parameters (Fig. 5 A and B).

#### The effect of fecundity and mortality

The success of eusocial groups depends on the ability of the reproductive individual to effectively use the resources it receives from members of the group. This ability is, however, critically limited by their maximum reproductive capacity *F_max_* which limits the amount of baseline mortality that can be compensated by reproduction. Thus, in population equilibrium, the group size of eusocial groups is limited to 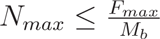, it increases with increasing fecundity *F_max_* (compare Fig. 5 H to E) and decreases with increasing mortality *M_b_* (compare Fig. 5 F to E). *F_max_* also influences the shape of the fertility function *(F*; Eqn. 5) and larger values of *F_max_* enlarge the convex part of this curve (Fig. 3 D). This increases the potential benefit eusocial groups can gain from increased inter-individual variance of resources for reproduction. Consequently, in eusocial groups resources needed in population equilibrium decrease and optimal group sizes severely increase with increasing values of *F_max_* (compare Fig. 5 D to A). A similar argument holds for increased cost of reproduction *c*_0_. *c*_0_ also increases the convex part of the fertility function (Fig. 3 C) and consequently increases the benefit of eusocial groups (Fig. 5 C).

For egalitarian groups the influence of *F_max_* on resources needed in population equilibrium as well as on optimal group sizes is far less pronounced. As larger values of *F_max_* enlarge the convex part of the fertility function and intra-specific competition reduces the amount of resources available for reproduction *x_r_*, the variance reducing effect of egalitarian resource sharing actually reduces mean fertility 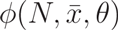. Thus, the amount of resources 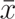 needed by egalitarian groups increases (compare Fig. 5 D to A) and group sizes of egalitarian groups decrease (compare Fig. 5 H to E) with increasing fecundity *F_max_*.

Baseline mortality *M_b_* has a similar effect on resource requirement. It increases the amount of resources 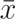 needed because higher baseline mortality must (in equilibrium) be compensated by higher reproduction and consequently by higher mean amounts of resources acquired. Costs of reproduction *C*0 on the other hand have only a negligible effect on the amount of resources needed 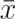 as well as on optimal group sizes *N_opt_* of egalitarian groups.

### Joining or leaving a group: evolutionary stability of eusocial groups

Whenever a large population of groups of size *N_pop_ >* 1 reaches a higher carrying capacity than a population of solitary individuals it can—in principle—not be invaded by individuals following a solitary strategy. In population equilibrium, the groups would drive mean resource availability below the critical value that allows the growth of a solitary strategy. This phenomenon is known from cooperatively breeding birds as “supersaturation” (Dickinson and Hatchwell, 2004).

However, this does not necessarily mean that groups in this population are evolutionarily stable and individuals have no incentive to leave them. In eusocial groups, subordinates do usually not reproduce. Their inclusive fitness is therefore determined solely by indirect fitness benefits, that is, the offspring of the related dominant individual (Hamilton, 1964a,b) if we ignore other direct benefits such as queueing for a dominant position (see, e.g., Kokko and Johnstone, 1999). Thus, it would clearly be beneficial to leave a group of size *N_pop_*—and such groups would become unstable—if the inclusive fitness of a solitary individual in a population of groups of size *N_pop_* exceeds that of a subordinate in a group of size *N_pop_*. In this light, the above described results hold true if relatedness equals one.

For our analysis of evolutionary stability of different strategies (i.e., different group sizes *N*), we use a simple fitness measure: the lifetime reproductive success of an individual (ψ). For our model lifetime reproductive success may be derived from mean fertility 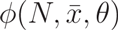 (Eqn. 7) and the mean lifetime 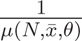 (according to Eqn. 4) of individuals as 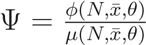. Here, ψ is a function of the size *N* of the group an individual is a member of, the mean size of resource items collected by individuals *θ* and the mean amount of resources collected 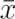. As the latter is itself an emergent property resulting from intra-specific competition in an equilibrium population of groups of size *N_pop_,* we may denote it as ψ(*N*, *N_pop_*, *θ*). In a habitat saturated by groups with a specific group size *N_pop_* = *N*, the rate of increase of groups of the same size will be ψ*(N_i_, N, θ)* ≠ 1 in equilibrium, while it may take different values ψ(*N*, *N*, *θ*) ≠ 1 for any other group size *N*_i_ when *N* ≠ *N_i_*.

So far we have used ψ as the mean fitness of a strategy. However, in eusocial groups the inclusive fitness ϕ of an individual depends on its role. The inclusive fitness ϕ_*sub*_(*N*, *N*, *θ*) of subordinates in a group of size *N* living in an infinitely large population of groups of the same size *N* is determined by their life expectancy 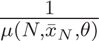 and the fertility 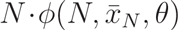 of the related dominant as

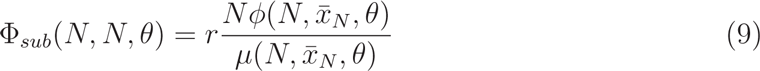

where *r* denotes the coefficient of relatedness while the mean amount of resources collected per individual 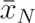 is a function of the population strategy *N*. When subordinates defect, leave the group and live as solitary individuals (*N_i_* = 1) they will loose indirect fitness benefits (as the related group now lacks one subordinate helper) but gain direct fitness benefits as a reproducing solitary individual. Now their inclusive fitness will be

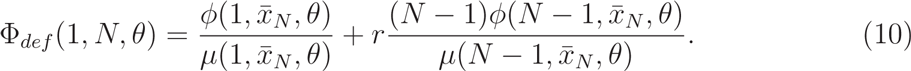

Note that, strictly speaking, this only holds as long as related individuals are not playing any evolutionary games (see, e.g., Hines and Smith, 1979). Subordinates should leave the group whenever leaving would result in a net increase in inclusive fitness, i.e when

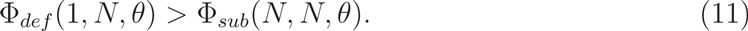

Eqn. 11 allows to derive the minimum relatedness *r_min_* preventing individuals from defecting, i.e. the minimum relatedness that allows the evolutionary stability of eusocial groups (see Fig. 6).

**Figure 6:**
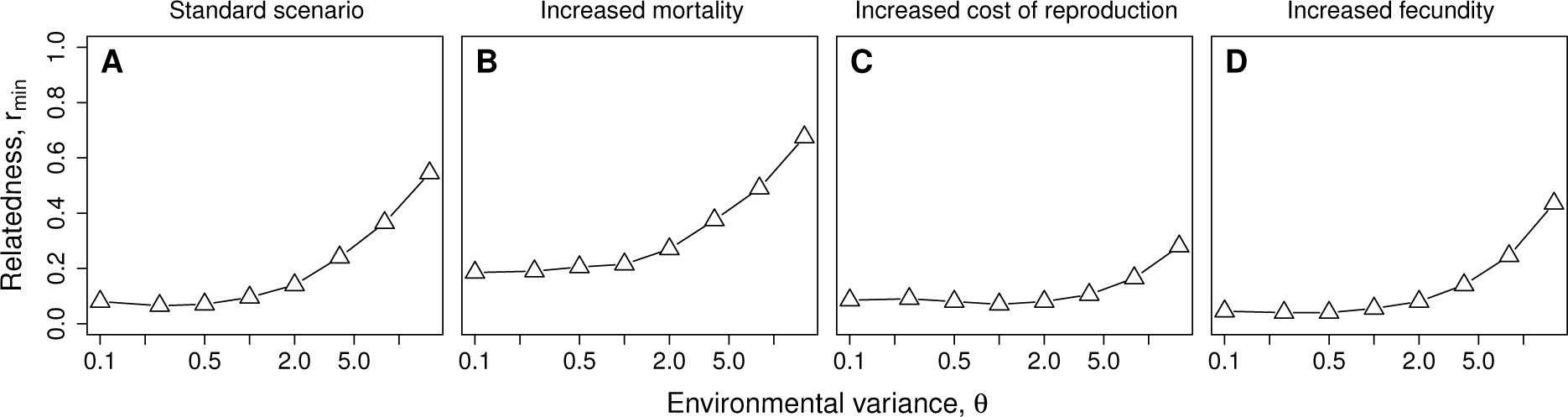
Influence of environmental variance (*θ*) on minimum relatedness *r_min_* of group members required to secure stability of despotic groups of optimal group size (*N_opt_*, see Fig. 5 E, F, G, and H) for four exemplary parameter combinations. (A): standard parameter set (*F_max_* = 3; *c*_0_ = 4; *M_b_* = 0.1; *o_M_* = 1); (B): increased mortality (*F_max_* = 3; *c*_0_ = 4; *M_b_* = 0.2; *o_M_* = 1); (C): increased cost of reproduction (*F_max_* = 3; *c*_0_ = 8; *M_b_* = 0.1; *o_M_* = 1); (D): increased fecundity (*F_max_* = 5; *c*_0_ = 4; *M_b_* = 0.1; *o_M_* = 1). Note the logarithmic x-axis.

Numeric solutions of Eqn. 11 (Fig. 6) show that particularly for low environmental variance *θ* the benefit of eusocial groups is sufficient to make the role of subordinate group members attractive even for individuals only modestly related to the dominant (*r_min_* ≈ 0.1). While increased baseline mortality *M_b_* (Fig. 6 B) increases the relatedness necessary for the stability of eusocial groups, increased cost of reproduction *c*_0_ (Fig. 6 C) and increased fecundity *F_max_* (Fig. 6 D) significantly decrease it and make eusocial groups evolutionarily stable even for extremely high environmental variance and low coefficients of relatedness (*r* < 0.25). The larger the optimal group size *N_opt_* and the larger the difference in mean resourced 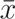 needed between solitary individuals and eusocial groups, the lower is the relatedness necessary to keep a subordinate individual in a eusocial group.

## Discussion

In contrast to previous work (e.g. Poethke and Liebig, 2008), the present model explicitly quantifies birth and death rates as functions of resource availability. This allows us to take into account competition for resources between individuals (see also Pen and Weissing, 2000) which reduces resource availability and ultimately results in *K*-selection. Furthermore, in comparison to Fronhofer et al. (2011a), the present study considers eusocial animal groups, that is, animal groups with reproductive skew. Our results show that—for a broad spectrum of model parameters—the formation of eusocial groups with resource sharing may severely increase carrying capacity (Fig. 5 A, B, C, and D). This will lead to the competitive exclusion of solitary foragers and breeders, a phenomenon known from cooperatively breeding birds as “supersaturation” (Dickinson and Hatchwell, 2004) and thus makes a reversion to solitary breeding less likely.

Our results demonstrate the potential of cooperative foraging and subsequent resource sharing (i.e. variance manipulation) as a driving force for the evolution of cooperative animal societies. Increased carrying capacity may thus have contributed to the evolution of eusocial animal groups. More importantly, this ecological benefit of group formation may have been important for the stabilization of cooperative breeding or eusociality after the transition from solitary life had already occurred, as our model does not explicitly consider the initial mechanism of group formation. Our model provides an ecological explanation for the benefit of group formation which sets it apart from previous models of reproductive skew (Vehrencamp, 1983; Reeve and Keller, 2001; Johnstone, 2000) that are often based on a predefined arbitrary benefit of group formation. In our simple consumer-resource model such an assumption is not required as group formation evolves because of the emergent advantages of variance manipulation.

As mentioned earlier, Poethke and Liebig (2008) demonstrate that egalitarian group formation, a variance reducing foraging strategy, is favored at high resource variances and that, by contrast, eusocial groups or cooperative breeding is advantageous when resource variance is low, because this strategy increases inter-individual variance in resource supply. However, when competition for resources is taken into account as in the present study, these predictions change. Eusocial groups remain at a clear advantage for low resource variances but become advantageous even for intermediate and rather high variance in resource availability (see Fig. 5). This is due to the twofold effect of eusocial groups on resource variance: inter-individual variance is indeed increased for reproduction, yet, for survival the opposite is true (Tab. 1).

### Model limitations

Throughout this work, we have analyzed the formation of eusocial groups under population equilibrium conditions. However, in a temporally and spatially heterogeneous landscape, and particularly in a metapopulation (Fronhofer et al., 2012), one will always find local populations that have not reached equilibrium density, yet. In newly colonized local habitat patches, for example, resources will be rather abundant and competition will be weak. This will necessarily favor solitary strategies with their high potential offspring numbers. Thus, landscape fragmentation and temporal heterogeneity in resource availability may lead to the coexistence of eusocial and solitary strategies.

While we do analyse the consequences of relatedness, and show that the ecological benefits of eusociality may be very large which makes eusocial groups evolutionarily stable even at low levels of relatedness, our modelling procedure implicitly assumes that groups have already been formed when we analyse the evolutionary stability of existing groups. Obviously, this may not be the case which limits the scope of our analyses and highlights that our model may be best thought as showcasing ecological benefits that are relevant for the maintenance and increase in size of already existing eusocial groups. Note that these restrictions do not apply to eusocial groups in which all members initially have a chance to become the dominant individual. Such groups can evolve by mutualism and indirect fitness benefits via relatedness are not be necessary (see e.g., Rissing et al., 1989).

A further limitation of our model is its comparison of only the two extreme cases of egalitarian versus eusocial groups, while in nature one will observe a continuum of cooperative strategies (see, e.g., Sherman et al., 1995). While this may impact our results quantitatively, the twofold benefit of forming eusocial groups discussed above remains a potentially important ecological mechanism responsible for the evolution and maintenance of eusocial groups. Of course, other factors (e.g., reviewed in Krause and Ruxton, 2002; Nowak, 2006; Lehmann and Keller, 2006) will also play a role for the evolution of eusociality and the relative importance of the different mechanisms may vary. Nevertheless, we suggest that our model is general in the sense that dealing with limited resources and variance in resource supply are challenges likely faced by a majority of organisms.

Finally, the ecological conditions considered here exclusively relate to the distribution and especially the variance in resource supply. While our model clearly shows the relevance of intraspecific competition for resources, we do not consider interspecific competition or predation, for instance (see Rankin et al., 2007; Tsuji, 2013).

### Empirical examples

It is interesting to note that, in our model, the increase in carrying capacity is generally more pronounced in eusocial than in egalitarian groups. Our model thus suggests that eusocial groups should dominate for a majority of environmental settings and life-history strategies. Although our model is very simple and compares only the extreme cases of egalitarian and eusocial groups, the dominance of eusocial groups in nature can be observed empirically: most cooperative societies are eusocial while truly egalitarian groups seem to be rare (Packer et al., 2001).

Typical eusocial groups are found among insects. In accordance with out model, the ubiquitously present and very successful ants alone show a bewildering array of different life-history strategies and feed on resources with typically low but also high variance (Hölldobler and Wilson, 1990). While these examples come from highly derived insect societies, our model may be more appropriate for primitively eusocial insects where subordinates are not sterile, for instance. An additional example are polistine wasps (reviewed in the context of skew theory in Reeve and Keller, 2001). While in the founding phase of a wasp nest the chance to become the reproductively dominant will make joining another female an attractive strategy, the probability to stay and accept the role of a “worker” will ultimately depend on the relatedness with the reproductively dominant individual. However, when an expensive nest is a prerequisite of successful reproduction this will change the shape of the fertility function. Such primary investments my be modeled as an offset that shifts the fertility function towards higher amounts of resources needed (Fronhofer et al., 2011a). Additional investments make reproduction more costly and will thus severely reduce the relatedness *r_min_* (see Fig. 6 C) necessary to stabilize despotic groups.

Cooperative systems with non-reproductive helpers can also be found in cooperatively breeding birds (Dickinson and Hatchwell, 2004) and the phenomenon of “supersaturation” has been well described in his context. In line with our results that predict an advantage of eusocial groups at low baseline mortalities, Arnold and Owens (1998) report that cooperatively breeding birds are generally characterized by low mortality rates. Furthermore, cooperative breeding seems to be consistently associated with low environmental variance in nature (Arnold and Owens, 1998, 1999; Ford et al., 1988; Gonzalez et al., 2013), although Jetz and Rubenstein (2011) find evidence for the opposite pattern. Our model corroborates these findings as it predicts an advantage for cooperative breeding and eusocial groups for both low and high resource variability.

By contrast, eusocial societies are rare in mammals (Clutton-Brock et al., 2009) where they have evolved only in four taxa: marmosets and tamarins, dogs, diurnal mongooses and African mole-rats. Typically, females in these groups show unusually high levels of fecundity.

Of course, also some examples of egalitarian groups are known. Social spiders have been discussed at length elsewhere (e.g. Fronhofer et al., 2011a). Our model predicts that egalitarian animal societies evolve when resource variance is high and offspring are few. These life-history traits are typically found in large mammals like lions (Packer et al., 2001) which do form egalitarian groups.

All theses examples show that global patterns of the occurrence of eusocial and cooperatively breeding groups—such as the influence of resource variance and life-history parameters (offspring cost and number)—in natural arthropod and vertebrate systems can, at least tentatively, be explained by the above presented model despite its great simplicity and caveats.

### Conclusions

In summary, pooling of resources for reproduction in eusocial groups severely increases intra-group variance in the amount of resources individuals may invest in reproduction. For upward convex fertility functions, eusocial groups thus out-compete solitary individuals as well as egalitarian groups. Whenever population growth is limited by resource availability, resources will necessarily be scarce and reproductive output will be dominantly determined by the convex part of the fertility function.

We thus show that competition for resources, i.e. density-dependence, and risk-sensitivity have the potential to lead to the evolution of cooperative breeding and eusociality (Fig. 5). More importantly, risk-sensitivity is likely important for the maintenance of eusocial groups and in the transition from small to larger groups that had previously formed due to other mechanisms. In our model, density-dependence is an emergent phenomenon and selection for increased resource-use efficiency leads to supersaturation (Dickinson and Hatchwell, 2004) of the environment, i.e. an increase in equilibrium density (Fig. 4).

Finally, our model yields some clear and testable predictions. In summary these are: (1) Eusocial groups are favored when offspring are numerous and cheap regardless of resource variance. (2) Egalitarian groups may evolve when resource variance is high and offspring are few and costly. (3) Increasing baseline mortality favors smaller eusocial groups and ultimately solitary living. (4) Eusocial groups can evolve and be maintained despite low levels of relatedness. (5) Globally, eusocial groups should be more frequent than egalitarian animal societies.

## Acknowledgements

The authors thank Hanna Kokko for comments on an earlier version of the text. E.A.F. is supported by Eawag. H.J.P. was supported by the German Research Foundation, DFG (SFB 554, TP C6).

